# Necrotizing enterocolitis causes increased ileal goblet cell loss in *Wnt2b* KO mice

**DOI:** 10.1101/2025.01.07.631715

**Authors:** Comfort Adegboye, Chidera Emeonye, Yu-Syuan Wu, Jaedeok Kwon, Luiz Fernando Silva Oliveira, Sathuwarman Raveeniraraj, Amy E. O’Connell

## Abstract

WNT2B is Wnt ligand which is able to support intestinal stem cells (ISC) in culture and support the intestinal epithelium in vivo. We have previously shown that WNT2B is critical for resistance to colitis, but not small intestinal injury, in the adult mouse. WNT2B is thought to coordinate with WNT3 in supporting ISC, and we have also shown that WNT3 expression is low in the early postnatal ileum in mice. Here, we hypothesized that WNT2B may be more critical in the small intestine during early development, and we challenged *Wnt2b* KO mice and controls with experimental necrotizing enterocolitis (NEC) on postnatal days 5-8. *Wnt2b* KO mice had similar ileum histology and injury scores to control mice. Molecular analyses showed that *Wnt2b* KO mice have differences in *Lgr5* and *Tlr4* expression compared to wild type controls in untreated conditions, but under experimental NEC expression of epithelial markers and inflammatory genes associated with NEC were similar to wild type. Periodic acid Schiff positive cells were lower in the villi of *Wnt2b* KO mice during NEC, however expression of goblet cell markers was not different compared to wild type mice. We also used an organoid-based NEC model to highlight the epithelium in isolation and also found no impact of WNT2B KO in the setting of NEC. These data further affirm that WNT2B is critical for inflammation responses in the mouse colon, but does not appear to play a major role in the small intestine, no matter the developmental period.

## INTRODUCTION

Intestinal Wnts are critical in the development and homeostasis of the epithelium ^1,2^. Our lab has shown that WNT2B is required for human intestinal health, as WNT2B loss-of-function (LOF) results in a novel type of congenital diarrheal enteropathy from the neonatal period ^3,4^. Subsequent studies from our lab have shown that WNT2B is particularly important in colonic homeostasis, as *Wnt2b* KO mice are significantly more susceptible to colitis compared to littermate controls ^5^. In the small intestine, however, the adult *Wnt2b* KO mouse appears to be more resilient than WNT2B LOF humans, as histology is normal at baseline and we saw no difference from controls after challenge with anti-CD3ε^5^. We have shown that there are regional difference in Wnt expression, with higher relative WNT3 in the small intestine, and higher WNT2B in the colon in mice ^5^. However, we have also shown that *Wnt2b* expression remains relatively stable throughout the first postnatal month in the mouse ileum, while *Wnt3* expression is low in the first week postnatally and increases over time^6^. Others have shown that Paneth cells, which are thought to be main contributors of WNT3 in the small intestine, confer resistance to NEC, and that mice are susceptible to NEC until Paneth cells fully develop around 2 weeks postnatal^7^. WNT2B and WNT3 have been shown to have functional redundancy in supporting intestinal epithelial stem cells and in maintaining epithelial homeostasis in vivo^1,8^. We therefore hypothesized that WNT2B may be more critical in the ileum in the early postnatal period, before total WNT3 increases.

Necrotizing enterocolitis (NEC) is an acute disease of the neonatal intestine that occurs mostly in premature infants, and which is frequently modeled using postnatal mice, owing to similarities in developmental features between mice and humans in these time periods^9^. Other groups have suggested that Wnt signaling may be important in the pathogenesis of NEC, as exogenous WNT7B was able to partially rescue the NEC phenotype in postnatal mice^10^. Given our interest in WNT2B, our knowledge that it is important in human small intestinal development, and its expression pattern over time in mice, we tested whether WNT2B LOF increases susceptibility to NEC in the immature intestine.

## Materials and Methods

### Research Ethics

All animal experiments were done in accordance with Boston Children’s Hospital Institutional Animal Care and Use Committee (IACUC, protocol 00001978). The study complied with ARRIVE guidelines.

### Mouse model

Wnt2b^fl/fl^ mice were a generous gift from T. Yamaguchi (NCI/NIH) ^11^. We bred the Wnt2b^fl/fl^ mice (C57Bl/6J background)with CMV-Cre mice (Jackson labs) to generate Wnt2b^fl/-^ mice, which were then crossed to generate Wnt2b KO mice. Male and female mice were used for experiments, and Wnt2b^+/+^ (wild type, WT) or Wnt2b^+/-^ heterozygous (het) matched littermates were used as controls. For all experiments WT and het mice were separately analyzed and no statistically significant differences were found between them, so they are presented as one group (WT) throughout the manuscript.

### Mouse model of Necrotizing Enterocolitis

We modified a previously published model of NEC^12^ for our protocol (Figure 1A). This is similar to other historical NEC protocols^13^. For our protocol, P5 pups were housed with the dam and removed to a 37°C warmer plate while they received a gavage with 28 kcal/oz neonatal formula plus 4mg/kg lipopolysaccharide (O111:B4, Sigma) via a trimmed 1.9Fr peripherally inserted central catheter line three times a day (8 hour intervals – 6am; 2pm; 10pm). Doses of formula increased from 50μL on treatment day one to 75μL on day 2 and 100μL on day 3. In addition, for two of these times points, the pups were also placed in a hypoxia chamber at 5% oxygen exposure for 10 minutes. This continued for 72 hours. Mice were weighed daily and examined for signs of illness or distress. Mouse NEC experiments were done in triplicate at a minimum, but due to slight unplanned variations between experiments, we did not group all the data into one output. For that reason some experiments have low n numbers, particularly in the control group. The results we report were consistent across all experiments.

**Figure 1.**
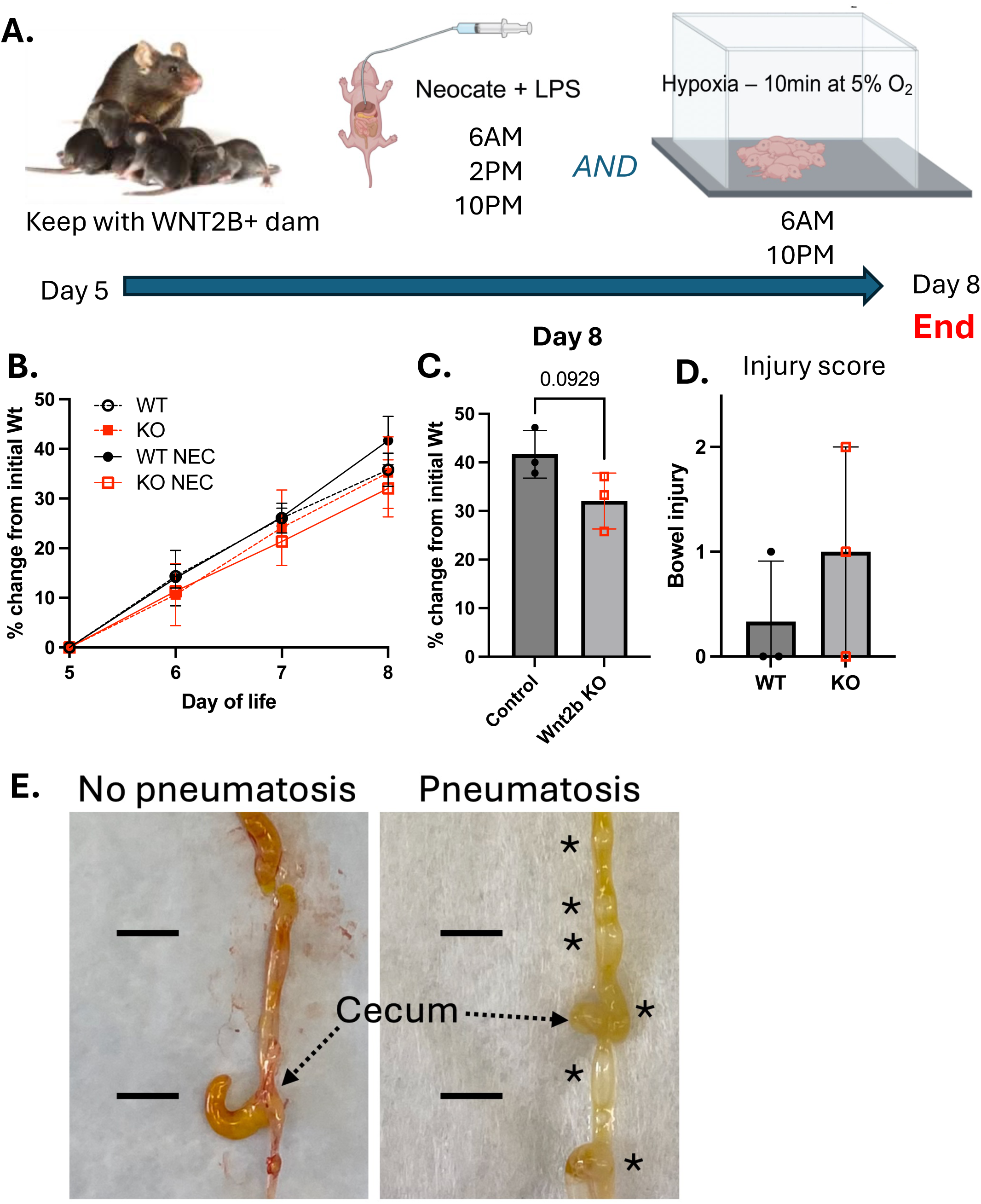
Gross and epithelial effects of NEC on *Wnt2b* KO mice. A.) Schematic of experimental approach. B.) Mouse weight as a percentage of the initial weight on the first day of treatment (P5). C.) Direct comparison of weight change between groups on day 8. D.) Gross injury score with 0 – normal, 1 - pneumatosis, 2 - bloody bowel. E.)

### Histological Analysis

Gross histologic assessment was performed to evaluate for evidence of pneumatosis intestinalis or necrosis at time of recovery. Grading was assigned in a blinded fashion with 0 for no evidence of injury, 1 for pneumatosis, and 2 for necrosis or perforation. For morphologic analysis, intestine or colon was cleaned of feculent material using 4% paraformaldehyde (PFA), and then tissue was sliced longitudinally, rolled and placed in a fixation cassette (Swiss roll method). Tissues were fixed in 4% PFA overnight at 4°C, then transitioned into 70% ethanol. Tissues were embedded in paraffin and sections (4-6μm) were stained with hematoxylin and eosin (H&E) or Periodic Acid Schiff (PAS). PAS+ cells were reported as the number of PAS+ cells per crypt/villous structure.

Scale bars are provided with details where possible; some of the microscopes we used did not provide integrated scale or were not functioning properly to integrate scale into the image, so magnification is indicated in the caption for these.

### Scoring for NEC

Slides were analyzed on an EVOS m7000 (ThermoFisher) at low magnification for imaging intact Swiss rolls (2x) or a Nikon Eclipse E800 microscope with Spot software for higher magnifications. Scoring for injury was done in a blinded fashion using a previously published scoring schema^14^: 0 - intact villi; 1 - superficial epithelial cell sloughing; 2 - mid-villous necrosis; 3 - complete villous necrosis; 4 - transmural necrosis. For colon analyses, we integrated inflammation scoring with a score of 0 for no inflammation, 1 for epithelial infiltration, 2 for submucosal infiltration, and 3 for muscular infiltration^5^.

### Quantitative PCR

Mouse tissue or HIOs were placed into Tri-reagent (ThermoFisher) and RNA was isolated using the Direct-zol kit (Zymo Research) or the RNAEasy mini kit (Qiagen) according to manufacturer’s instructions. RNA was then reversed transcribed into cDNA using a high-capacity cDNA reverse transcription kit and RNAse Inhibitor (ThermoFisher) according to manufacturer’s instructions. Quantitative PCR was performed on a QuantStudio6 Flex (ThermoFisher) using Taqman qPCR Master mix and specific primers. Taqman primers used in analyses are indicated in Supplemental Table 1.

### Mouse intestinal enteroid and colonoid generation

Mouse organoids were generated as described previously^15–18^. Intestinal tissue, either duodenal or colonic, was cleaned using cold intraluminal phosphate buffered saline (PBS), cut into smaller pieces, and cleaned in PBS until the supernatant was clear. The tissue was then placed in 2mM EDTA (small intestine) or 5mM EDTA (colon) and incubated on ice with rocking for 30 minutes. The tissue was then shaken vigorously for 2 minutes to release crypts and pipetted up and down 25 times with a 10mL serological pipet while mixing. The solution was filtered and centrifuged. Dissociated cells were then washed in Advanced DMEM/F12 (Gibco) and reconstituted in Matrigel (Corning, 50μL/well) plated in 24-well plates and incubated for 10 minutes at 37°C. Next, 0.5mL growth media was added to the wells (Table 1). Enteroids were fed with EGF/Noggin/R-spondin (ENR) growth media every 2-3 days and passaged every 7 days.

### In vitro NEC assay

Organoids were treated with LPS 0.1mg/mL in ENR media and then placed in a hypoxia chamber at 5% oxygen content (wash out with nitrogen) according to a previously published protocol for in vitro NEC^10^. After 24 hours, organoids were imaged and then collected for analyses. This experiment was repeated in duplicate with similar results in each experiment.

## RESULTS

### Wnt2b KO mice are affected by NEC similarly to control wild type littermates

From P5-P8, mice were gavage-fed three times a day with hyperosmolar formula plus LPS (Figure 1A). Twice daily, they were also subjected to hypoxia exposure in a 5% oxygen chamber for 10 minutes, immediately following a feeding. Mice were kept with the dam otherwise to ensure adequate hydration and support. *Wnt2b* KO mice tended to gain less weight by day 8, but this was not statistically significant (Fig. 1B, 1C). Injury on gross bowel examination was similar to wild type controls (Fig. 1D). Microscopic histology with hematoxylin and eosin staining showed swollen epithelium in NEC-treated mice, but this appeared similar between *Wnt2b* KO mice and controls (Fig. 1E). Histologic epithelial injury was also similar to controls using the maximum injury throughout the intestine (Fig. 1F), and the proportion of the intestinal epithelium that showed injury was also similar to control wild type mice (Fig. 1G).

### NEC causes decreased epithelial PAS positive cells in Wnt2b KO mice compared to controls

We also stained epithelium with Periodic Acid Schiff (PAS), which stains protein rich structures pink. In the epithelium this is most usually goblet cell granules in the villi and Paneth cell granules at the base of the crypt (Fig. 2A). As we would expect at this age, there were not organized Paneth cell granules in the crypts in any of the conditions. Granules from goblet cells stained positive, and notably there were significantly less PAS positive cells in the *Wnt2b* KO mice treated with NEC (Fig. 2B). Other groups have identified loss of goblet cells as a hallmark of human NEC biopsy tissues^19–21^.

**Figure 2.**
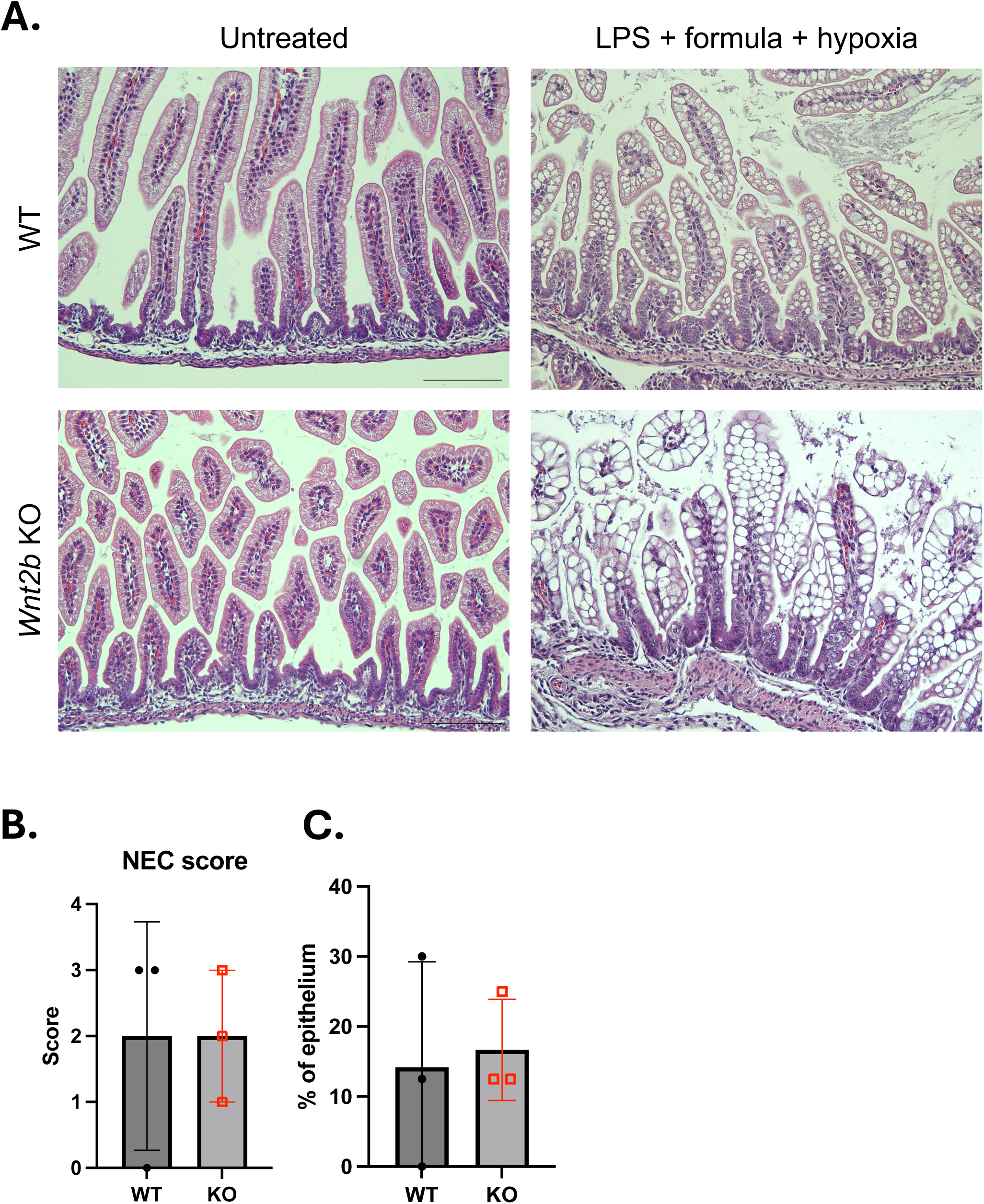
Epithelial injury in *Wnt2b* KO mice and wild type controls after NEC. A.) H&E staining of ileum. Scale bar = 200μm. B.) Epithelial NEC score. C.) Percentage of bowel with injury.

### Wnt2b KO mice have RNA differences at baseline but similar expression to controls in the setting of experimental NEC

Given the histologic findings, and our previous work showing that WNT2B loss-of-function causes decreased ISC markers, we did qRT-PCR for key intestinal epithelial lineage markers and for inflammation genes previously implicated in NEC^22–24^ (Fig. 3A.) *Lgr5* was significantly decreased in untreated *Wnt2b* KO mice in the ileum. Expression of *Lyz1*, a Paneth cell marker, was lower in NEC treated control mice, but higher in Wnt2b KO mice compared to controls in untreated and treated conditions, although this did not reach significance and the fold change was not large. *Tlr4* expression was surprisingly decreased in our NEC treated animals, and *Wnt2b* KO mice had lower baseline expression of *Tlr4* compared to controls. *Cxcl10*, which has been shown to decrease in experimental NEC, trended to be lower in NEC-treated control mice, but was unchanged in the *Wnt2b* KO mice. In alignment with the finding of fewer PAS+ goblet cells in the *Wnt2b* KO mice, *Muc2* expression was also decreased in the *Wnt2b* KO mice, which was consistent with the lower number of PAS+ cells on histology.

**Figure 3.**
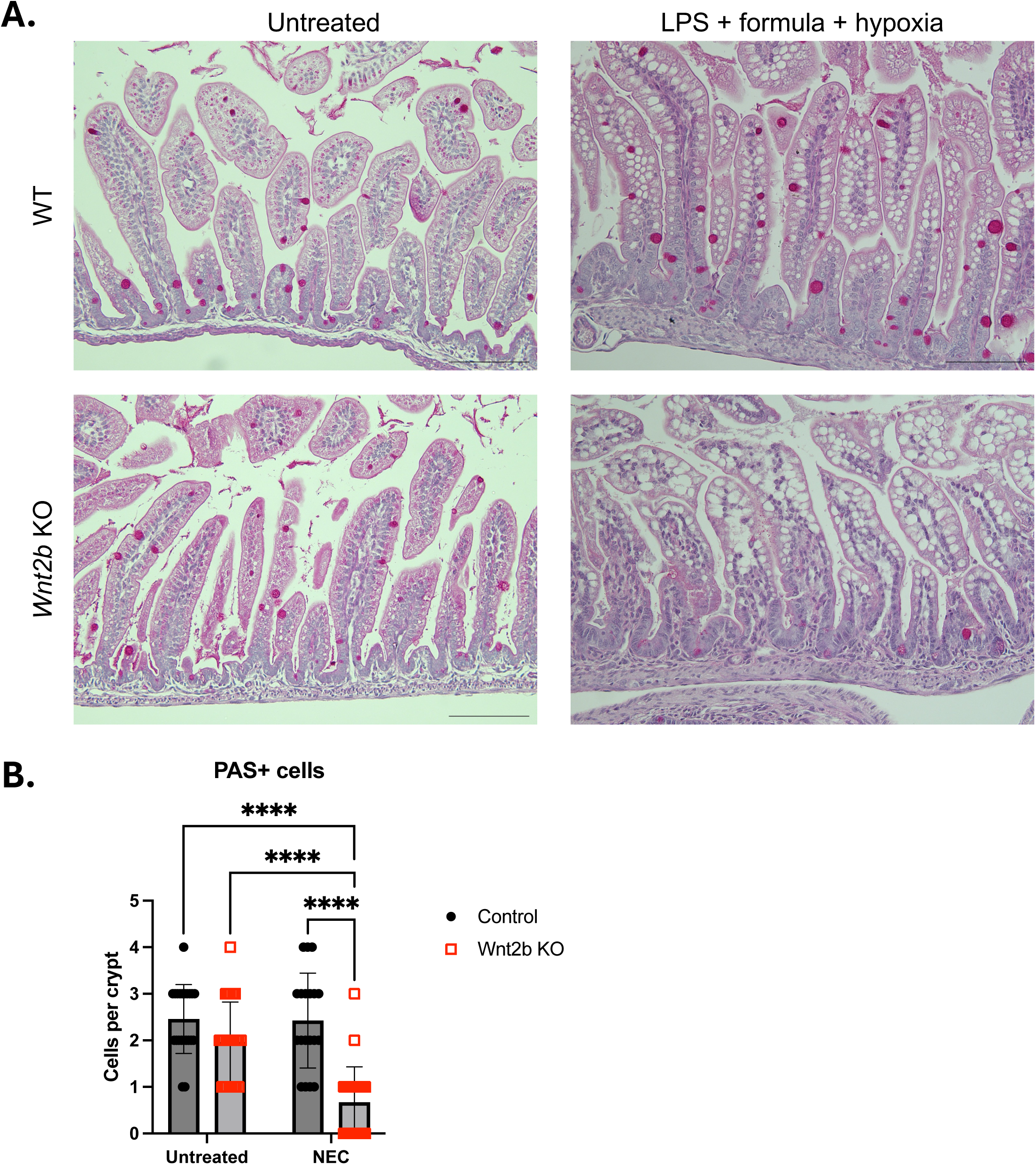
Periodic acid Schiff staining in *Wnt2b* KO mice and wild type controls after NEC. A.) PAS stained epithelium from untreated and NEC treated mice at P8. Scale bar = 200μm. B.) Manual count of PAS-positive cells per crypt/villus unit. **** p < 0.0001.

### Wnt2b KO mice did not have increased impacts from NEC in the colon

Because WNT2B is important for preventing inflammation in the mature colon^5^, and because the mouse model of NEC can impact the colon, we evaluated the impacts of these experiments on colon tissue. Gross histology was similar between *Wnt2b* KO mice and control mice treated with NEC (Fig. 4A). We did note significant immune cell infiltration in the colon, but this was similar between *Wnt2b* KO and control NEC-treated mice. In contrast to the ileum, goblet cell numbers were similar between *Wnt2b* KO and control mice as assessed on PAS staining (Fig. 4B). Epithelial damage, inflammation, and muscle thickening in the colon were also similar between groups (Fig. 4C). Expression of *Lgr5* was significantly lower in the *Wnt2b* KO mouse colon after NEC, but expression of *Lyz1, Muc2, Tlr4*, and *IL-6* was similar between *Wnt2b* KO mice and controls after NEC.

**Figure 4.**
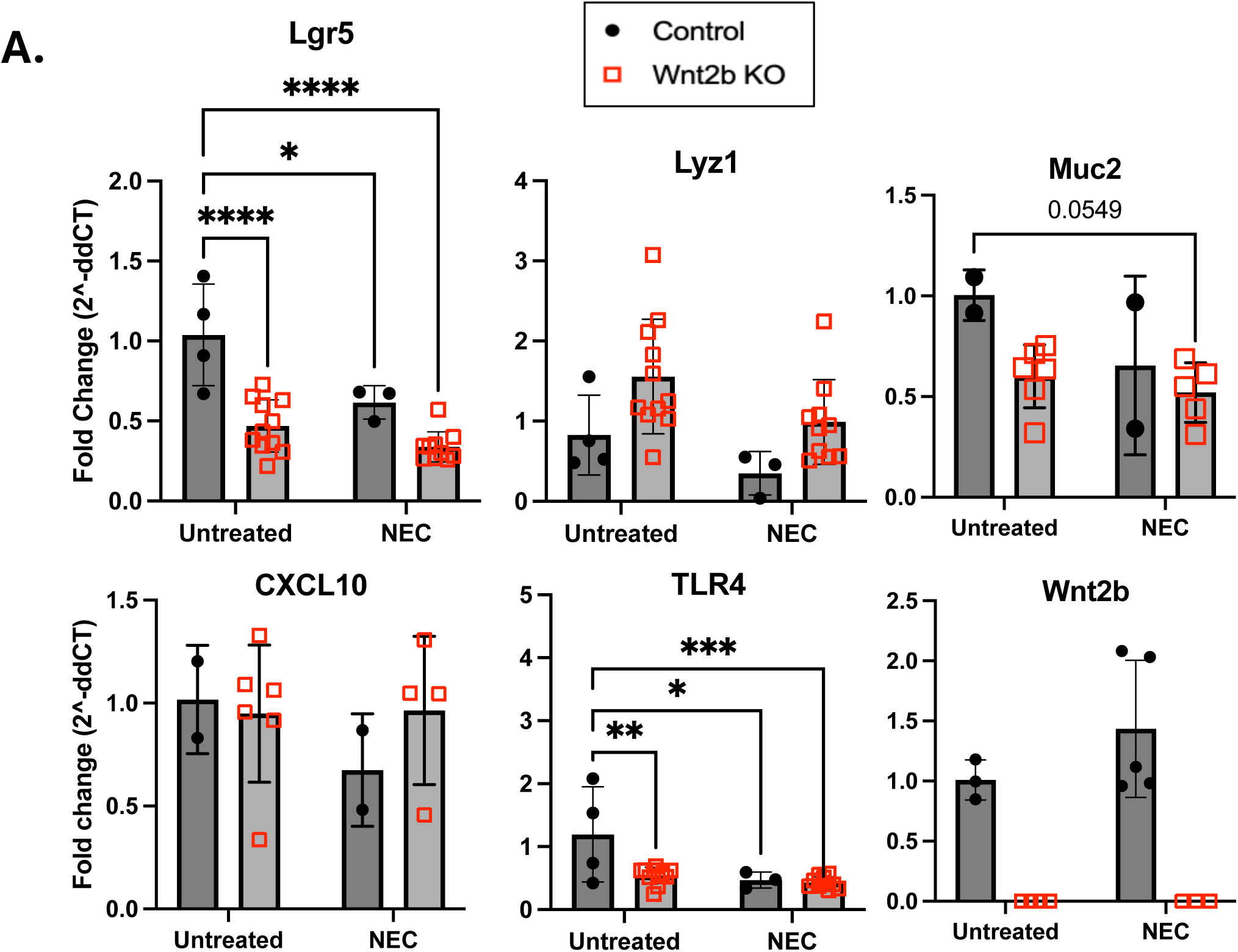
Expression of epithelial lineage markers and inflammatory genes by qRT-PCR. A.) qRT-PCR comparison of RNA expression between wild type and Wnt2b KO mice at day 8 in untreated controls and NEC mice. * p < 0.05, ** p < 0.01, *** p < 0.001, **** p < 0.0001.

### Breastmilk did not contribute to the effects of Wnt2b on NEC phenotype

Proteomic analyses have shown that breastmilk contains WNT2B^25^. We therefore repeated the NEC experiments using a *Wnt2b* KO dam (Fig. 5A), in order to ensure that the mice were not receiving WNT2B in the breastmilk during the experiment, since they were kept with the dam and allowed to breastfeed ad lib. Weights during the experiment were similar between *Wnt2b* knock out mice and controls (Fig. 5B). Histology also showed similar mild injury in both *Wnt2b* KO mice and controls (Fig. 5C). Epithelial damage scores were not different between groups (Fig. 5D). The *Wnt2b* KO mice appeared grossly to have less dense villi after NEC in this experiment, so we counted the number of villi per unit of length and did not detect any difference between groups (Fig. 5D). Similar to experiments using wild type dams, *Wnt2b* KO mice treated with NEC had significantly fewer PAS positive cells (Fig. 5E, 5F).

**Figure 5.**
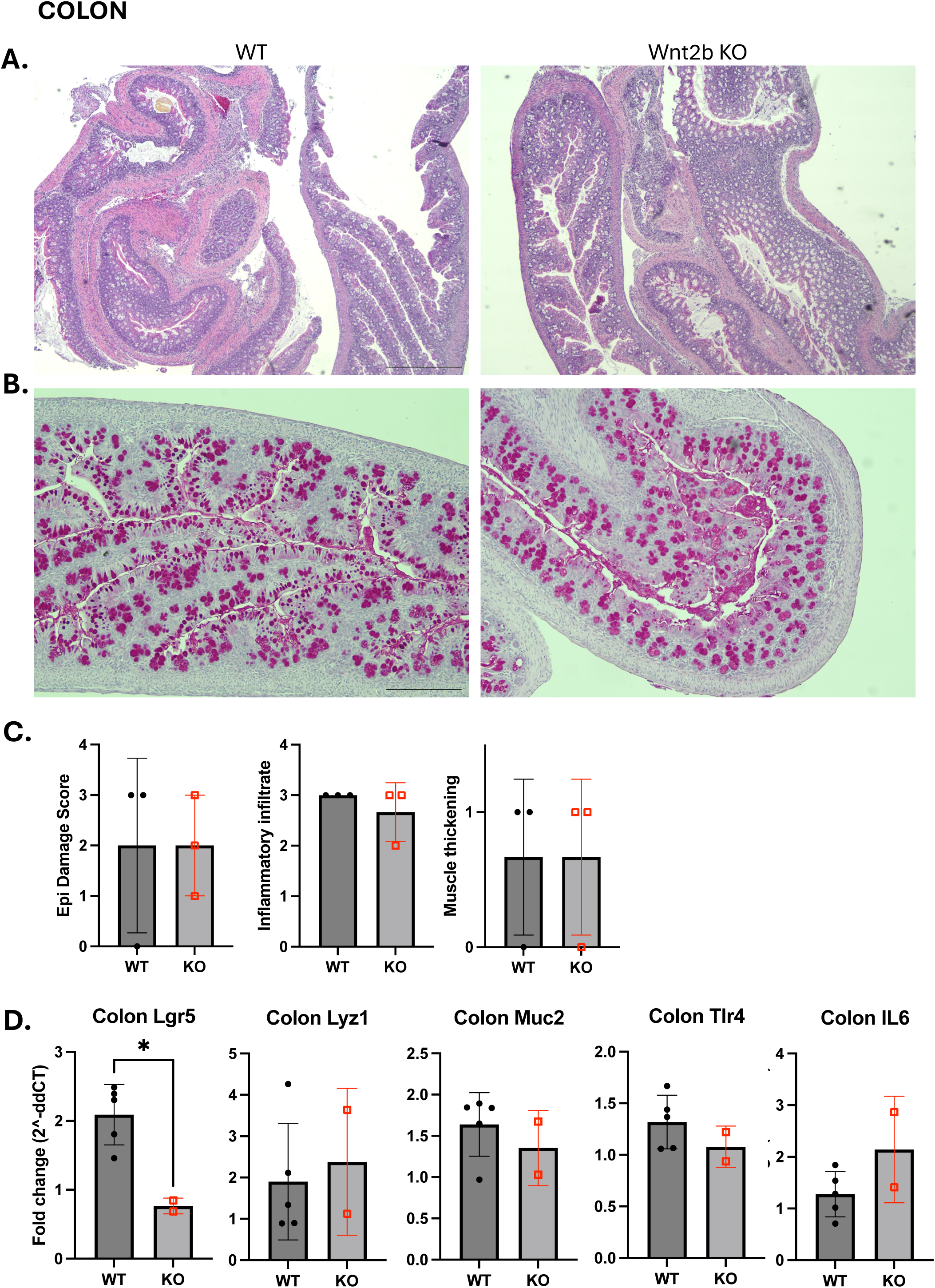
Colon histology and gene expression after NEC. A.) H&E staining comparing wild type and Wnt2b KO mice after NEC. 4X magnification. B.) PAS staining showing goblet cells at 10X magnification. C.) Colon epithelial damage scores and inflammation scores. D.) Gene expression by qRT-PCR. * p < 0.05

### Wnt2b KO epithelial organoids

WNT2B is primarily produced by the intestinal mesenchyme, but can also be produced autonomously in the epithelium^5,26^. To scrutinize the contribution of epithelial WNT2B in protection against early small intestinal injury, we isolated ileum organoids from the ileums of 7-day old *Wnt2b* KO and wild type littermate mice. We then exposed the organoids to LPS and hypoxia according to previously published methods for inducing NEC in vitro^10^ (Fig. 6A). Both wild type (WT) and *Wnt2b* KO organoids appeared darker than untreated organoids, indicating increased luminal debris, but otherwise there were no morphologic differences in the organoids (Fig. 6B). qRT-PCR analyses of the organoids again showed decreased *Lgr5* in *Wnt2b* KO organoids at baseline, and lower expression in NEC conditions but no difference in the KO mice versus WT. *Lyz1* expression trended downward in NEC-treated organoids, as did *Muc2* in the *Wnt2b* KO mice. Experimental NEC again decreased expression *Tlr4*, as well as *Tnfa*, while *IL-6* expression was significantly induced in the WT organoids but not *Wnt2b* KO organoids after NEC (it was not expressed or expressed at very low levels in untreated organoids).

**Figure 6.**
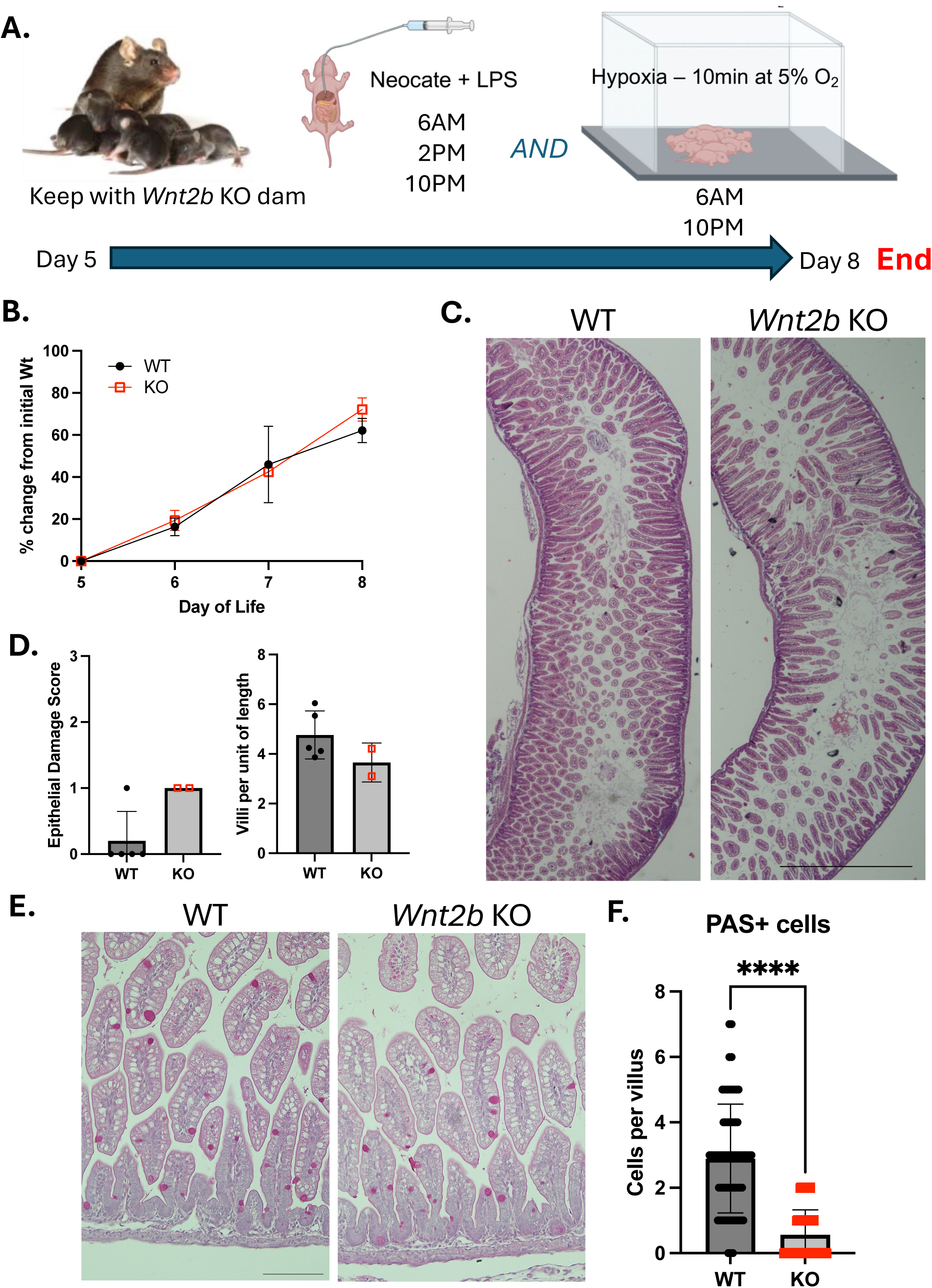
Gross and epithelial effects of NEC on control and *Wnt2b* KO mice when fed by *Wnt2b* KO dam. A.) Schematic of experimental approach. B.) Mouse weight as a percentage of the initial weight on the first day of treatment (P5). C.) H&E staining of ileum at 4X. D.) Epithelial NEC score and density of villi. E.) PAS stained epithelium from untreated and NEC treated mice at P8. Scale bar = 200μm. F.) Manual count of PAS-positive cells per crypt/villus unit. **** p < 0.0001.

**Figure 7.**
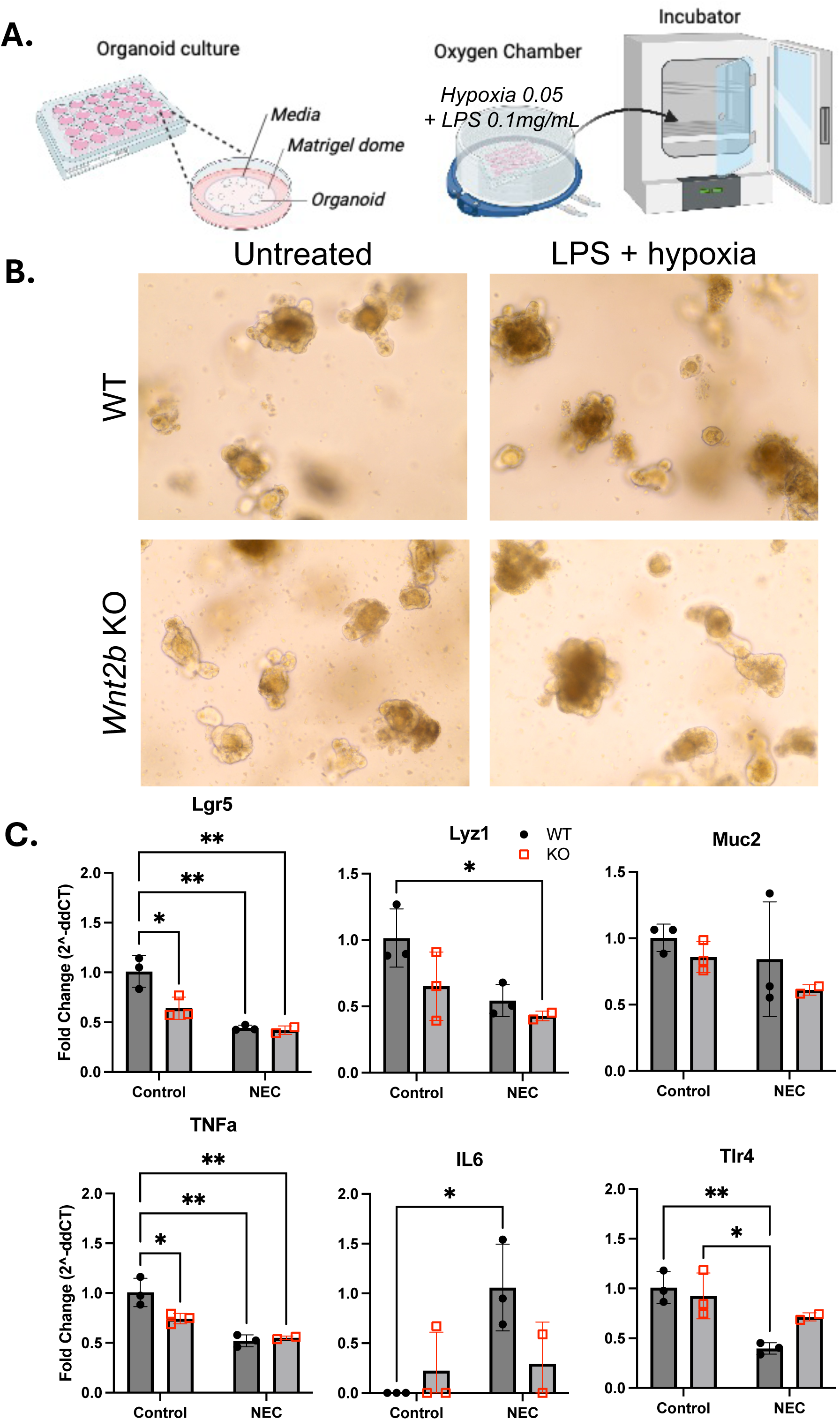
*Wnt2b* KO and control epithelial organoids treated with NEC in vitro. A.) Schematic of experimental approach. B.) Appearance of organoids at the end of the experiment. 10X magnification on EVOS XL. C.) Gene expression by qRT-PCR. * p < 0.05, ** p < 0.01.

## DISCUSSION

Here we investigated whether WNT2B is needed for early life protection against inflammation in the small intestine using an experimental model of NEC. We hypothesized that it may have higher importance in early development, since expression of *Wnt3* is lower than in mature mice in this period while *Wnt2b* expression is stable over time^6^. The *Wnt2b* KO mice demonstrated a mildly increased phenotype compared to controls, with more loss of goblet cells according to PAS staining and expression of *Muc2*. Overall this is a mild phenotype, indicating that loss of WNT2B is not catastrophic in the small intestine in mice. Wnt signaling has been previously implicated in NEC pathogenesis by other groups. One in vivo NEC study examined Wnt signaling in the context of the already injured intestine and found that NEC was associated with decreased expression of *Wnt3a*, *Wnt5a*, and *Wnt7a*, while *Wnt3* levels were increased in NEC tissue^10^ (did not remark on WNT2B). This post-NEC analysis of expression, however, may be more of a signature of cellular injury and may not be particularly specific to NEC nor related to the pathogenesis. The authors then treated mice with recombinant WNT7B before experimental NEC and showed partial amelioration of the phenotype^27^. *Wnt7b* expression was decreased in murine NEC compared to baseline in their RNA analysis. *Wnt7b* was not detected in our RNA analysis of the small intestine of C57Bl/6J mice over the first postnatal month^6^.

The decrease in PAS+ cells in the *Wnt2b* KO mice support a mild phenotype, where there is some early injury that is consistent with human NEC, which also shows diminished goblet cells^10,19,21^. Others have shown that production of TNFα in the setting of NEC may trigger mucin loss from goblet cells in neonatal mice^21^. In our in vitro analysis TNFα was decreased in the NEC conditions, so it is unclear if the mucin loss in the *Wnt2b* KO mice was via the same mechanism.

*Lgr5* expression was decreased in *Wnt2b* KO mice, and treating with NEC conditions reduced control *Lgr5* expression to similar levels. *Wnt2b* KO mice did not experience any further decrease in *Lgr5* expression with NEC, however. This suggests that the residual ISC that are able to be maintained in the absence of WNT2B are resistant to further injury via the LPS plus hypoxia model. Alternately, it is possible that new ISC are being generated sufficiently to keep up with turnover after injury.

Interestingly, the colon in *Wnt2b* KO mice was not more affected than wild type mice in the NEC model. This is in contrast to dextran sodium sulfate (DSS) colitis in mature (2 month) mice, where we saw significantly increased susceptibility to injury in *Wnt2b* KO mice^5^. We did see evidence of colitis in the mice, with inflammatory cell infiltrates. It is likely that the chemical disruption caused by DSS is more potent in epithelial disruption than the NEC model we used. Indeed, the NEC model is used because it is thought to approximate human NEC, which most often affects the ileum. The colon also had normal numbers of goblet cells in the *Wnt2b* KO mice, in contrast to the ileum, suggesting more resistance to loss of goblet cells in the colon than the ileum.

WNT2B can be detected in human breastmilk^25^, so we repeated the experiment using *Wnt2b* KO dams to ensure there was no WNT2B in the mothers’ milk. We saw no increase in phenotype with this, again there was goblet cell depletion but similar overall injury to controls. We also performed an in vitro NEC assay using epithelial-only organoids to evaluate the effects of loss of WNT2B in the epithelium devoid of mesenchymal and endothelial cells. We again saw that *Lgr5* expression was lower in the absence of WNT2B, but *Muc2* expression was not significantly different from controls. This may be due to organoid media containing exogenous Wnts that skew the population toward stem cells. We saw that *TNFα* expression was decreased in the NEC model, as was *Tlr4*, however control organoids demonstrated an increase in *Il6* expression compared to baseline.

We used a published model of NEC that was reported to result in up to 50% animal death by 3 days of treatment in C57Bl/6J mice. We did not experience this degree of death in our protocol. This may be due to slight variations in commercial LPS strain. The mice did develop pneumatosis and epithelial injury, although the epithelial injury was limited to patches and not diffuse. The phenotype we observed was consistent with NEC (pneumatosis, some epithelial sloughing, loss of ISC and goblet cells), making us confident the approach was effective. Since we hypothesized that *Wnt2b* KO mice would have more severe injury with NEC, it was tolerable that our experimental phenotype was not severe; if our phenotype in wild type mice was profound we would not be able to observe increased injury in the knock out mice.

In conclusion, WNT2B confers some resistance to injury in experimental NEC. *Wnt2b* KO mice demonstrated mildly increased injury with significant loss of goblet cells. Additionally, WNT2B does contribute to some resistance to injury in the small intestine in early development, at least in mice.

## Abbreviations

EGF: epidermal growth factor
GI: gastrointestinal
KO: knockout
Lgr: leucine-rich repeat-containing G-protein coupled receptor
NEC: necrotizing enterocolitis
SI: small intestine
Wnt: wingless-related integration site protein
WT: wild type

## Author Contributions

All authors read and critically reviewed the manuscript and approved of the final manuscript. In addition:

CA performed all organoid experiments and contributed to mouse experiments.

CE managed the mouse colony and contributed to mouse experiments.

SW performed experiments and managed the mouse colony.

JK, LFO and SR assisted with in vivo experiments and some experimental analyses, as well as experimental planning.

AEO conceptualized the project, conceptualized and supervised all of the experiments, and wrote/edited the manuscript.

## Data Availability Statement

All raw data is available upon reasonable request by emailing the corresponding author.

